# High-throughput prediction of peptide structural conformations

**DOI:** 10.1101/2024.12.03.626727

**Authors:** Alexander M. Ille, Christopher Markosian, Stephen K. Burley, Paul C. Whitford, José N. Onuchic, Wadih Arap, Renata Pasqualini

## Abstract

Protein structure prediction *via* artificial intelligence/machine learning (AI/ML) approaches has sparked substantial research interest in structural biology. In recent work, AlphaFold2 (AF2) has been adapted for the prediction of multiple structural conformations, beyond static structures, by using multiple random seeds and subsampling the multiple sequence alignment (MSA). Research applying this specialized approach has focused on regular-sized proteins, while smaller peptides remain underexplored in this context. Here, we report AF2-based multi-conformation prediction and systematic analyses of a set of 557 peptides (10–40 residues in length) with corresponding nuclear magnetic resonance (NMR) ensemble coordinates. We found that predicted ensembles varied in similarity to experimental coordinate data, with disulfide-rich peptides bearing somewhat closer resemblance whereas α-helical peptides exhibited greater dissimilarity. Distinct correlations involving MSA depth were also observed. These results reveal notable capabilities of AF2-based multi-conformation prediction for peptides but also highlight considerable limitations, emphasizing the need for interpretation discretion and improved multi-conformation prediction approaches.

## INTRODUCTION

AlphaFold2 (AF2) is an artificial intelligence/machine learning (AI/ML) model capable of predicting the three-dimensional (3D) structures of proteins from amino acid sequence alone with accuracy comparable to lower-resolution experimentally determined protein structures [1, 2]. AF2 was trained on multiple sequence alignment (MSA) of metagenomic sequencing data in combination with protein structures determined by X-ray crystallography and cryo-electron microscopy (cryoEM) from the Protein Data Bank (PDB) [3]. Since its release, AF2 has garnered widespread use for various applications across the biological sciences [4], such as modeling large protein complexes [5-7] and proteome-scale structure prediction [8, 9]. More recently, AF2 has been adapted for predicting the structures of multiple protein conformations, going beyond the original scope of single static structure prediction [10-13]. This is accomplished by using multiple random seeds, *i*.*e*., stochastic prediction initializations, and subsampling of the input MSA, resulting in an ensemble of predicted structures [12]. MSA subsampling, in particular, introduces co-evolutionary sequence-based variability by dedicating unique subsets of the MSA for different structures in a predicted conformational ensemble. The prediction of multiple conformations with AF2 has been demonstrated to align with experimentally determined conformational ensemble coordinate data for certain proteins as determined by nuclear magnetic resonance (NMR) spectroscopy [10, 11], which is notable considering that the training data for AF2 did not contain NMR structures [1, 14]. However, AF2-based multi-conformation prediction is a distinct application for which accuracy is expected to vary, as with the prediction of static structures, and should therefore be considered with discretion [12]. Furthermore, the focus of such research has been on regular-sized proteins, while prediction of structural conformations for peptides in this context remains underexplored.

Prediction of peptide structures is of great relevance for elucidating biological processes as well as for the development of peptide-based therapeutics, and AF2 has been a subject of ongoing research and evaluation for application towards these purposes [14, 15]. A previous study by McDonald et al. [16] benchmarked the prediction capabilities of the conventional AF2 approach for a rich dataset of 588 peptides with NMR-determined 3D structures in the PDB. The conventional AF2 approach was originally intended for static structure prediction and computes five structures for a given input sequence [1, 12]. In contrast, the aforementioned multi-conformation AF2 approach involving multiple seeds and MSA subsampling computes a total of 80 structures per prediction run [12]. The study by McDonald et al. [16] utilized the conventional AF2 approach [1], which differs in scope and intended application from the multi-conformation AF2 approach [10-13], the latter of which has not yet been systematically explored in the context of peptides. The inherent conformational variability of peptides merits further consideration if AI/ML approaches are to be utilized for the discovery and characterization of peptide function and peptide-protein interactions, both of which are structurally dynamic in nature. Advances in this domain are anticipated to facilitate experimental research involving peptides in contexts ranging from biomedical to industrial, including *in vivo* phage display for the discovery of therapeutically relevant peptides and their target receptors [17-22], the design of peptides for agricultural disease control and sustainability [23-26], and peptide-based biomaterial development [27, 28], among others.

Herein, we report AF2-based multi-conformation prediction of peptide structures by using multiple random seeds and MSA subsampling. High-throughput prediction of conformational ensembles was performed for *n* = 557 peptides from the McDonald et al. [16] dataset with *n* = 80 individual structures computed per peptide, providing a combined total of 44,560 individual structures. Computational analyses involving various metrics, including root-mean-square deviation (RMSD), root-mean-square fluctuation (RMSF), and MSA sequence depth, were used to compare predicted and experimental structures. Our analyses revealed that AF2-based conformational ensemble prediction of peptides varied in similarity to NMR coordinate data, with a median RMSD of 1.63 Å and a median RMSF difference of 0.688 Å across the entire set of 557 peptides. Notably, predictions of disulfide-rich peptides tended to exhibit closer agreement with NMR ensemble coordinates in terms of conformational variability, particularly relative to α-helical peptides. Moreover, we compared the multi-conformation AF2 approach with the conventional AF2 approach and found that certain peptides more closely resembled NMR coordinate data, depending on the utilized approach. The current work provides an assessment of the experimental consistency of peptide conformational ensemble prediction, expanding upon the scope of conventional AF2 prediction. While these results highlight notable capabilities of this multi-conformation prediction approach, substantial limitations are also present, underscoring the need for discretion in prediction interpretation and improved prediction methodologies.

## RESULTS

### Prediction of peptide conformational ensembles and MSA depth associations

AF2 predictions rely on the model’s pre-trained weights as well as the MSA generated for a given input sequence [1, 12]. With the central role of MSA in AF2 predictions, we first asked whether MSA depth had systematic impacts on the accuracy of peptide ensemble predictions. In addition, we further explored whether these impacts are consistent across the peptide groups. To predict conformational ensembles using the AF2 multi-conformation approach [10-13] **(Fig. 1a)**, we utilized the ColabFold implementation of AF2 which provides a specialized workflow for introducing structural variability *via* random seed initializations and MSA subsampling [12]. The number of seeds was set to 16, with five structures per seed (*n* = 80 structures per prediction), and MSA subsampling parameters were set to 16 for number of sequence cluster points and 32 for extra sequences (see Methods for additional information). An example of a conformational ensemble prediction and its assessment is shown in **Fig. 1b-c** for an antimicrobial peptide derived from *Amaranthus caudatus* (PDB 1ZUV) [29]. The example highlights an area of conformational variability **(Fig. 1b)** and depicts per-residue RMSF differences (ΔRMSF) *versus* the NMR conformational ensemble **(Fig. 1c)**. The original dataset by McDonald et al. [16], which contained 588 peptides, was filtered to include only peptides for which the NMR ensemble contained five or more structural conformers in order to provide conformational variability while avoiding substantial reduction in sample size **(Fig. 1d)**. Peptide categorizations from the original dataset were adopted in the current study in order to assess how conformational prediction similarity to experimental structures might vary among these groupings, which include α-helical membrane-associated peptides (AH MP), α-helical soluble peptides (AH SL), β-hairpin peptides (BHPIN), disulfide-rich peptides (DSRP), and mixed secondary structure peptides (MIX). Accordingly, *de novo* structural predictions were performed for a total of *n* = 557 peptides, consisting of *n* = 172 AH MP peptides, *n* = 37 AH SL peptides, *n* = 56 BHPIN peptides, *n* = 261 DSRP peptides, and *n* = 31 MIX peptides **(Supplementary Table 1)**.

**Fig. 1:**
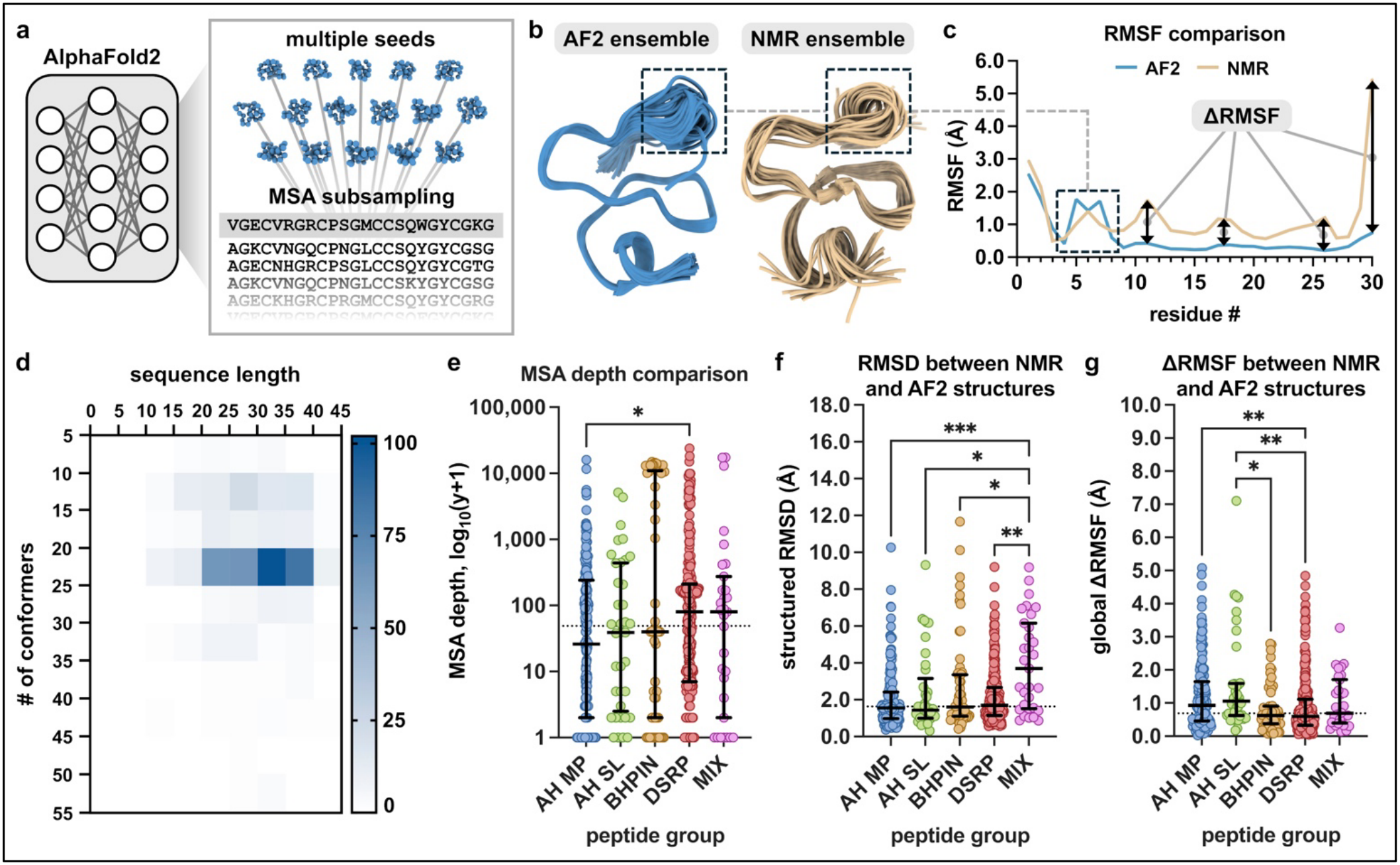
Peptide structural conformation prediction and analysis. **(a)** Overview of the AF2-based multiple conformation prediction approach with multiple seeds and MSA subsampling. **(b)** AF2-predicted and NMR-determined conformational ensembles of an antimicrobial peptide (PDB 1ZUV). A flexible region is outlined with a dashed box. **(c)** Comparison of the Cα RMSF values from the conformational ensembles presented in (b). A flexible region present in both ensembles is outlined with a dashed box. Examples of individual ΔRMSF values are also indicated with arrows. **(d)** Heatmap of the *n* = 557 peptide NMR dataset displaying sequence length *versus* number of conformational structures per ensemble, with the scale bar representing the number of PDB entries. **(e)** MSA depth (number of sequences within the MSA) for each peptide among the six different peptide groupings. Data represented as medians and corresponding interquartile ranges (IQRs), with the dotted line indicating median across all peptides. Statistical analyses by Kruskal Wallis one-way ANOVA with post-hoc Dunn’s multiple comparisons test, \**p* = 0.0171. To accommodate the wide and uneven distribution of the data, the y-axis was plotted on a log_10_ scale following addition of 1 to all values, *i*.*e*., log_10_(y+1), allowing for inclusion of MSAs with values of zero. Statistical analyses were performed using untransformed values (prior to log transformation). **(f)** Structured region Cα RMSD between AF2 and NMR conformational ensemble coordinates among the six peptide groupings. Data are represented as medians ± IQRs, with the dotted line indicating median across all peptides. Statistical analysis by Kruskal Wallis one-way ANOVA with post-hoc Dunn’s multiple comparisons test, \**p* = 0.0104 (AH SL *vs*. MIX), \**p* = 0.0446 (BHPIN *vs*. MIX), \*\**p* = 0.0059, \*\*\**p* = 0.0001. **(g)** Cα global ΔRMSF **(eqn. 1)** between AF2 and NMR conformational ensemble coordinates among the six peptide groupings. Data are represented as medians ± IQRs, with the dotted line indicating median across all peptides. Statistical analysis by Kruskal Wallis one-way ANOVA with post-hoc Dunn’s multiple comparisons test, \**p* = 0.0195, \*\**p* = 0.0027 (AH MP *vs*. DSRP), \*\**p* = 0.0022 (AH SL *vs*. DSRP). For panels e-g, peptide groupings include the following: AH MP *n* = 172 peptides; AH SL *n* = 37 peptides; BHPIN *n* = 56 peptides; DSRP *n* = 261 peptides; MIX *n* = 31 peptides.

The MSAs generated by ColabFold (*via* MMseqs2) [12, 30] had a median depth of 48 sequences across all peptides with large distributions among the different peptide groups, and the DSRP group had a modestly greater MSA depth than the AH MP group **(Fig. 1e)**. RMSD analysis of alpha carbons (Cα) of structured regions [16] between the AF2-predicted and NMR-determined conformational ensemble coordinates was performed for each individual peptide (see Methods). The median of the structured region RMSD between predicted and NMR ensemble coordinates across all 557 peptides was 1.63 Å, and among the distinct peptide groups only the MIX group exhibited statistically different (greater) structured region RMSD **(Fig. 1f)**. To quantify the similarity in conformational variability between the predicted and NMR conformational ensemble coordinates at corresponding residues, RMSF differences were assessed using a global ΔRMSF metric **(eqn. 1)**.

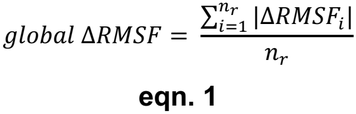

where *i* is the residue position, *n*_*r*_ is the number of residues, and ΔRMSF **(Fig. 1c)** is the per-residue difference between the individual RMSF values of two conformational ensembles, with each ΔRMSF taken as an absolute value (see Methods).

The median of the global ΔRMSF was found to be 0.688 Å across all peptides. The DSRP group was found to have a lower global ΔRMSF compared to the AH MP and AH SL groups, while the BHPIN group also had a lower global ΔRMSF compared to the AH SL group **(Fig. 1g)**. A possible explanation for the DSRP group observation is that disulfide bonds generally promote the formation of well-defined conformations, potentially making them simpler prediction targets. When considering all peptides (ungrouped), there was no significant correlation between amino acid sequence length and global ΔRMSF **(Fig. S2a)**. However, evaluating the peptide groups separately revealed weak positive correlations for the AH MP group (Spearman *r* = 0.3570, *p* < 0.0001) **(Fig. S2b)** and AH SL group (Spearman *r* = 0.3724, *p* = 0.0232) **(Fig. S2c)**. This indicates a tendency toward reduced similarity between predicted and NMR ensemble coordinates as sequence length increases for α-helical peptides. The BHPIN, DSRP, and MIX groups did not exhibit significant correlations between sequence length and global ΔRMSF **(Fig. S2d-f)**. Comparisons of average RMSF values of individual ensembles, as opposed to global ΔRMSF between ensembles, further indicate that flexibility of α-helical peptides (AH MP and AH SL) was substantially underestimated for AF2 predictions compared to NMR references **(Fig. S3a and b)**, while predictions for the DSRP group were only marginally underestimated **(Fig. S3d)** and no significant differences were observed among the BHPIN and MIX groups **(Fig. S3c and e)**.

It is notable that the DSRP group had both a greater MSA depth and a lower global ΔRMSF compared to the AH MP group in the independent assessment of these variables **(Fig. 1e and g)**, given that AF2 leverages co-evolutionary signals from MSAs. We further considered whether any broader relationship was present between MSA depth and global ΔRMSF **(Fig. 2)**. Although there was no significant correlation found across the entire set of peptides **(Fig. 2a)**, assessment of the different protein groups revealed mixed trends, with some groups exhibiting distinct correlations **(Fig. 2b-f)**. The DSRP group exhibited a weak negative correlation (Spearman *r* = -0.2670, *p* < 0.0001) **(Fig. 2e)**, where greater MSA depth may contribute to slightly lower global ΔRMSF between predicted and experimental ensembles. Paradoxically, the AH MP and AH SL groups, both of which are predominantly α-helical, exhibited weak positive correlations (AH MP Spearman *r* = 0.2771, *p* = 0.0002; AH SL Spearman *r* = 0.3400, *p* = 0.0395) **(Fig. 2b and c)**. Although these correlations are weak, it is worth noting that the inconsistency of positive correlations among the AH MP and AH SL groups *versus* the negative correlation within the DSRP group aligns with the global ΔRMSF comparisons **(Fig. 1g)**, where DSRP peptides exhibit modestly greater similarity between predicted and experimental ensembles than both AH MP and AH SL peptides. In summary, these analyses reveal weak but distinct correlations between MSA depth and agreement between predicted and experimental conformational ensembles among certain peptide groups, despite the absence of an overall correlation.

**Fig. 2:**
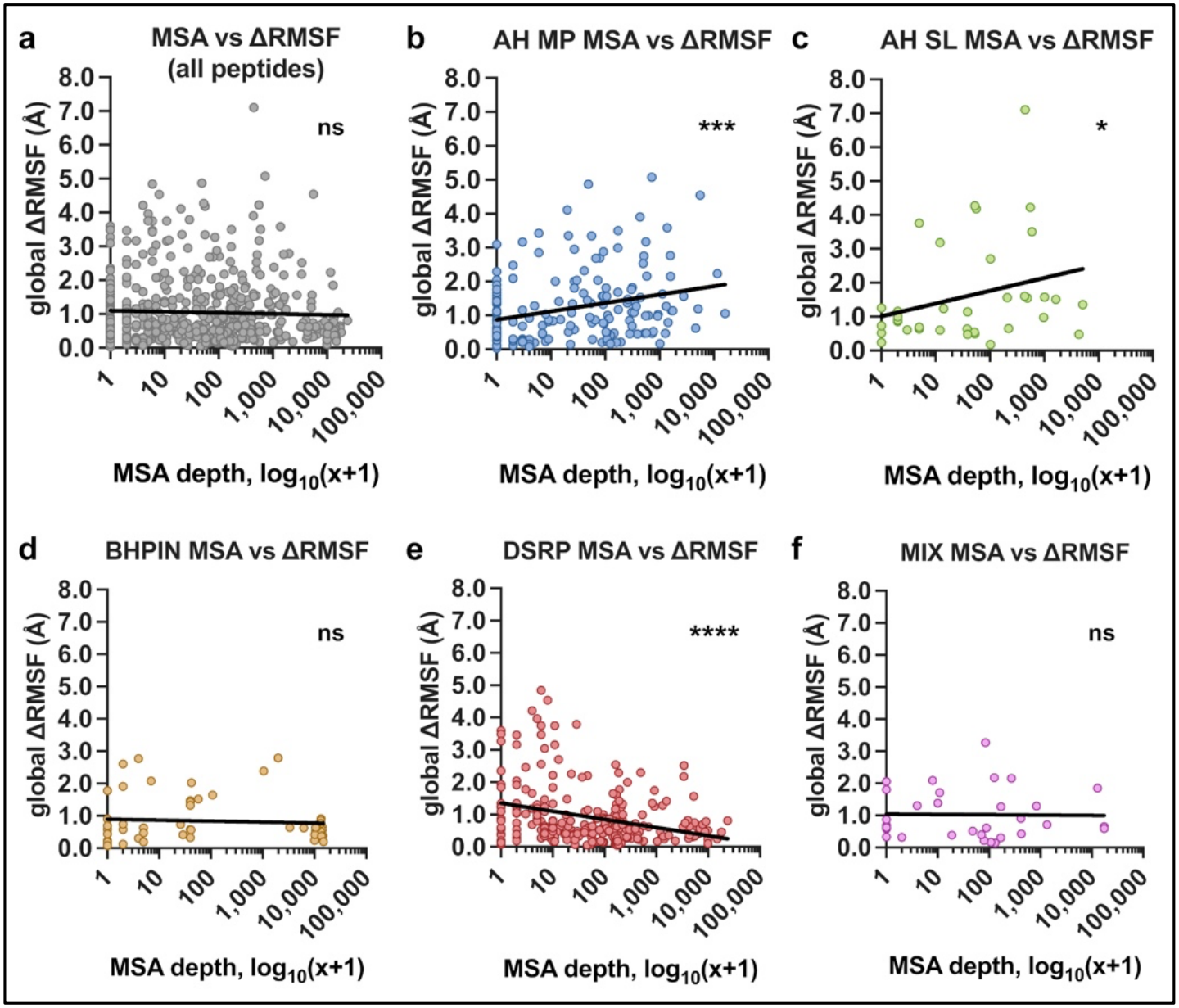
Associations of MSA depth with global ΔRMSF between AF2-predicted and NMR-determined structural ensembles. **(a-f)** MSA depth *versus* Cα global ΔRMSF for all peptides (A) and for the different groups (B-F). Statistical analyses by Spearman correlation: **(a)** all peptides *r* = -0.0174, 95% CI -0.1028 to 0.0682, ns (not significant, *p* ≥ 0.05), *n* = 557 peptides; **(b)** AH MP *r* = 0.2771, 95% CI 0.1286 to 0.4134, \*\*\**p* = 0.0002, *n* = 172 peptides; **(c)** AH SL *r* = 0.3400, 95% CI 0.0081 to 0.6045, \**p* = 0.0395, *n* = 37 peptides; **(d)** BHPIN *r* = 0.0919, 95% CI -0.1829 to 0.3535, ns, *n* = 56 peptides; **(e)** DSRP *r* = -0.2670, 95% CI -0.3793 to -0.1469, \*\*\*\**p* < 0.0001, *n* = 261 peptides; **(f)** MIX *r* = -0.0073, 95% CI -0.3702 to 0.3575, ns, *n* = 31 peptides. For all panels, the x-axes were plotted on a log_10_ scale following addition of 1 to all values, *i*.*e*., log_10_(x+1), allowing for graphical representation of wide and uneven MSA distributions and inclusion of MSAs with values of zero. Statistical analyses were performed using untransformed values (prior to log transformation). Trend lines (depicted in black) were generated by nonlinear regression using a semilog line fit (logarithmic x-axis, linear y-axis).

### Conformational variability and example peptides

We next considered whether the trends in global ΔRMSF could be rationalized in terms of the degree of variability present in each NMR ensemble. That is, some peptides may exhibit greater conformational variability, whereas others may be more structurally constrained. Additionally, NMR-determined structural ensemble coordinates reflect non-uniform and method-dependent criteria for inclusion/exclusion of conformers, which can influence the apparent conformational variability of deposited ensembles [31, 32]. In order to sort peptides in a manner that considers both conformational similarity and the intrinsic variability of the NMR reference, we used a composite score **(eqn. 2)** calculated as the ratio of two quantities: (i) the global ΔRMSF between AF2-predicted and NMR-determined ensemble coordinates; and (ii) the average RMSF of the NMR ensemble coordinates (RMSF_NMR_). The composite score ranks peptides by prioritizing similarity between predicted and NMR ensemble coordinates while weighting by the magnitude of NMR ensemble coordinate variability, such that lower scores indicate ensembles with increased similarity along with greater experimental flexibility **(Fig. 3a; Supplementary Table 2)**.

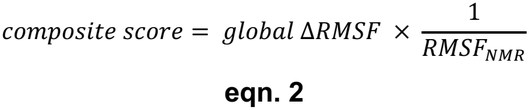

**Fig. 3:**
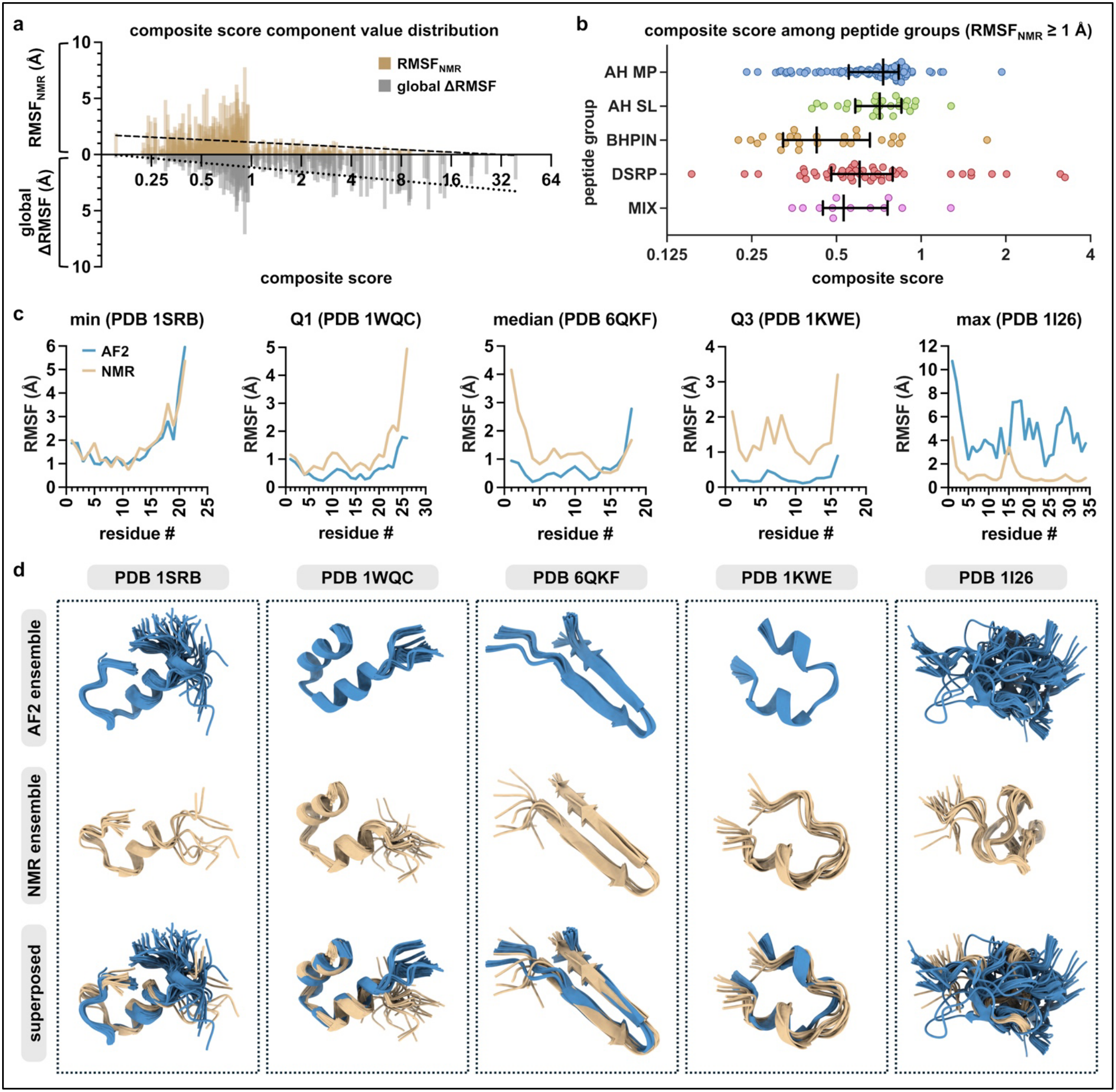
Analysis of conformational ensemble flexibility and quartile-representative peptide examples. **(a)** Bidirectional plot of the distribution of the composite score for each conformational ensemble (x-axis) along with corresponding Cα RMSF_NMR_ (upward y-axis) and Cα global ΔRMSF (downward y-axis) component values of the composite score. To accommodate the wide and uneven distribution of the composite score values, the x-axis was plotted on a log_2_ scale. Trend lines were generated by nonlinear regression using a semilog line fit (logarithmic x-axis, linear y-axis), with the dashed line representing RMSF_NMR_ values and the dotted line representing global ΔRMSF values. **(b)** Distribution of the composite score for each conformational ensemble among the different peptide groups including only ensembles with RMSF_NMR_ values ≥ 1.0 Å, consisting of: AH MP *n* = 104 peptides; AH SL *n* = 26 peptides; BHPIN *n* = 25 peptides; DSRP *n* = 47 peptides; MIX *n* = 12 peptides. Data represented as medians and corresponding IQRs. **(c)** Comparisons of Cα RMSF values of AF2 and NMR conformational ensemble coordinates representative of the minimum, Q1, median, Q3, and maximum composite scores from the DSRP peptide group for peptides with RMSF_NMR_ ≥ 1.0 Å. Since quartile values are interpolated and may fall between observed values, representative ensembles were selected as the largest observed composite score ≤ each quartile. **(d)** AF2-predicted (blue), NMR-determined (beige), and superposed structures of the representative conformational ensembles from (c). Global ΔRMSF values are as follows: PDB 1SRB, 0.2920 Å; PDB 1WQC, 0.6020 Å; PDB 6QKF, 0.8060 Å; PDB 1KWE, 1.0943 Å; and PDB 1I26, 3.5995 Å.

where *global ΔRMSF* is the overall RMSF difference between two conformational ensembles, as defined in **eqn. 1**, and *RMSF*_*NMR*_ is the corresponding average RMSF of the NMR ensemble.

However, the composite score is also sensitive to peptides that have very low ΔRMSF despite having limited flexibility within the NMR ensemble, *i*.*e*., sub-angstrom RMSF_NMR_, thereby obscuring identification of peptides which exhibit both high conformational similarity and high flexibility **(Fig. S4)**. Accordingly, further analyses were focused on peptides with RMSF_NMR_ ≥ 1.0 Å among each of the five peptide groups **(Fig. 3b)**. Quartile-representative peptides for the minimum, Q1, median, Q3, and maximum composite score for each peptide group vary substantially in terms of similarity between predicted and NMR ensemble coordinates **(Fig. 3c and d; Fig. S5-8)**. As exemplified within the DSRP group, per-residue RMSF comparisons between predicted and NMR ensemble coordinates of peptides with lower composite scores tend to exhibit better agreement, while poor agreement is increasingly evident with higher scores, particularly beyond the median **(Fig. 3c and d)**. This general tendency is also present in the other peptide groups **(Fig. S5-8)** and aligns with trends of increasing global ΔRMSF across composite score quartile groupings **(Fig. S9)**. Together, these data suggest that accounting for the degree of experimental conformational variability may provide useful context for distinguishing peptides with differing levels of agreement between predicted and experimental ensembles.

### Comparison of multi-conformation and conventional AF2 approaches

We further asked how the multi-conformation AF2 approach and conventional AF2 approach compare with respect to experimental ensemble similarity and if the multi-conformation approach is more suitable in some context. We therefore compared the two approaches in terms of agreement to NMR ensemble coordinates as indicated by global ΔRMSF **(Supplementary Table 3)**. The conventional AF2 approach uses a single seed which generates an output of five individual structures per prediction, allowing for calculation of global ΔRMSF in a similar manner (though with fewer structures) to the multi-conformation AF2 approach, which uses 16 seeds and generates an output of 80 individual structures. Pairwise comparisons of ensembles predicted by the multi-conformation approach and the conventional approach were performed across all peptides and among the different peptide groups **(Fig. 4a)**, as well as restricted to peptides with RMSF_NMR_ ≥ 1.0 Å **(Fig. 4b)**. Across the entire set of peptides, there was no significant difference found between the global ΔRMSF of ensembles predicted by the two separate approaches, regardless of whether the RMSF_NMR_ ≥ 1.0 Å criteria was applied. Within the AH MP group, ensembles predicted by the multi-conformation approach had statistically greater global ΔRMSF than those predicted by the conventional approach. The DSRP group predictions did not exhibit any significant difference between approaches for all peptides within the group. However, for DSRP peptides with RMSF_NMR_ ≥ 1.0 Å, global ΔRMSF was significantly lower for the multi-conformation approach compared to the conventional approach. The MIX group exhibited a marginally significant difference, but not for peptides with RMSF_NMR_ ≥ 1.0 Å, whereas no significant differences were found among the AH SL and BHPIN groups.

**Fig. 4:**
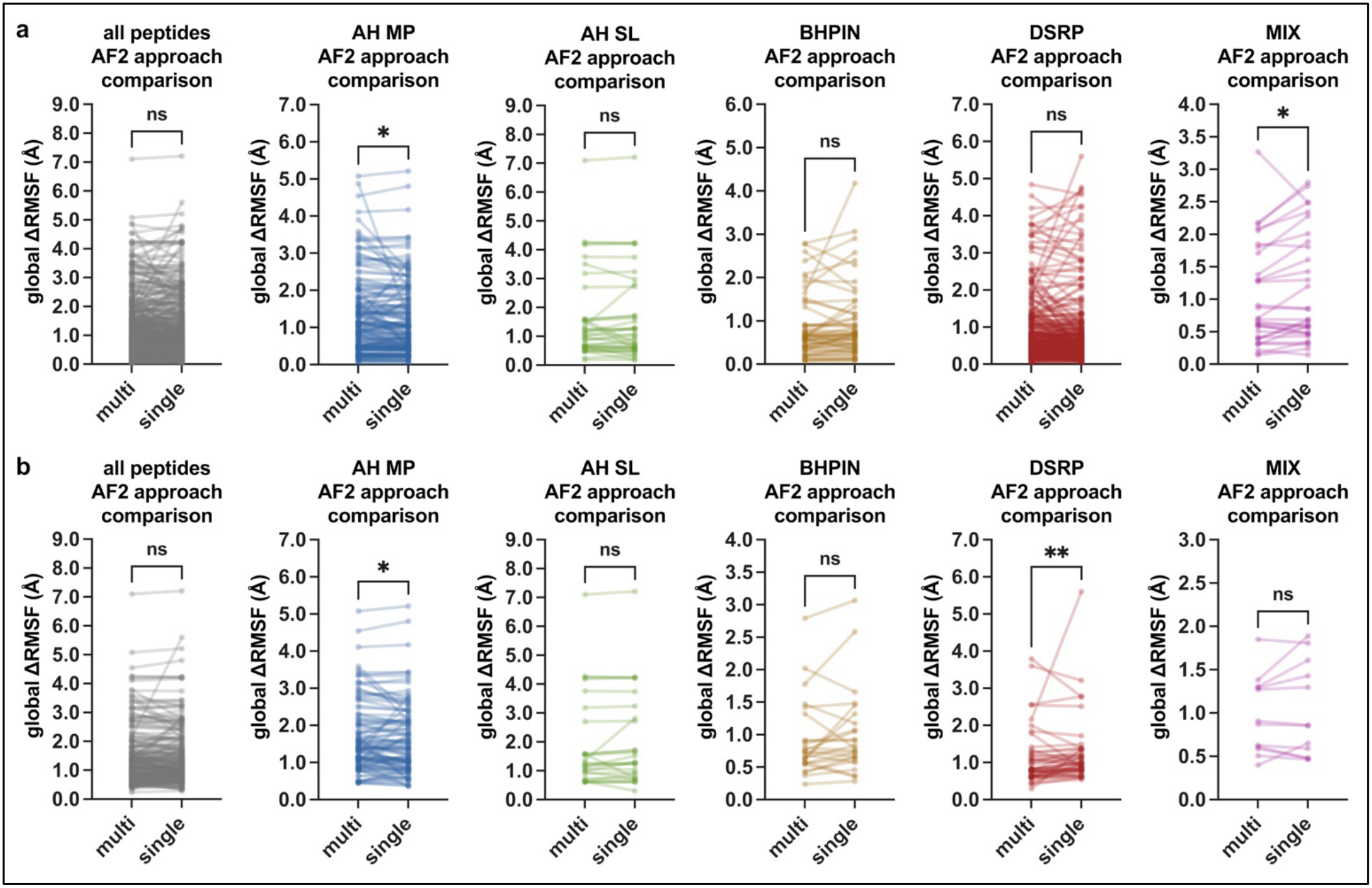
Comparison between the multi-conformation and conventional AF2 approaches. Global ΔRMSF between predicted and NMR ensemble coordinates for the multi-conformation AF2 approach using multiple seeds and MSA subsampling (multi) and for the conventional AF2 approach using single seeds without subsampling (single). Connecting lines indicate corresponding peptides from each prediction approach. **(a)** Comparisons of all 557 peptides together (grey) and by peptide groupings (colored), including: AH MP *n* = 172 peptides; AH SL *n* = 37 peptides; BHPIN *n* = 56 peptides; DSRP *n* = 261 peptides; MIX *n* = 31 peptides. Statistical analysis by Wilcoxon matched-pairs test, AH MP \**p* = 0.0219 (median of multi group = 0.9312 Å, median of single group = 0.8543 Å), MIX *p = 0.0374 (median of multi group = 0.6917 Å, median of single group = 0.6867 Å), ns (not significant, p ≥ 0.05). **(b)** Comparisons of peptides with RMSF_NMR_ ≥ 1.0 Å together (grey) and by peptide groupings (colored), including: AH MP *n* = 104 peptides; AH SL *n* = 26 peptides; BHPIN *n* = 25 peptides; DSRP *n* = 47 peptides; MIX *n* = 12 peptides. Statistical analysis by Wilcoxon matched-pairs test, AH MP \**p* = 0.0357 (median of multi group = 1.3926 Å, median of single group = 1.3934 Å), DSRP \*\**p* = 0.0077 (median of multi group = 0.8121 Å, median of single group = 0.9749 Å), ns (not significant, p ≥ 0.05).

In addition to the pairwise statistical comparisons, we used categorical percentages to summarize which prediction approach resulted in greater similarity to NMR coordinate variability, *i*.*e*., lower global ΔRMSF **(Fig. 5)**. Across the entire set of peptides, ∼50% of ensembles predicted by the multi-conformation approach had lower global ΔRMSF compared to the conventional approach, regardless of whether the criteria of RMSF_NMR_ ≥ 1.0 Å was applied **(Fig. 5a)**. For the AH MP group, only 38.3% (no inclusion/exclusion criteria) and 35.6% (RMSF_NMR_ ≥ 1.0 Å) of ensembles predicted by the multi-conformation approach had lower global ΔRMSF compared to the conventional approach **(Fig. 5b)**. The AH SL, BHPIN, and DSRP groups all had more ensembles with lower global ΔRMSF predicted by the multi-conformation approach, and to a greater extent for peptides with RMSF_NMR_ ≥ 1.0 Å **(Fig. 5b-e)**. Notably, the greatest percentage (70.2%) of ensembles with lower global ΔRMSF predicted by the multi-conformation approach was found to be for peptides with RMSF_NMR_ ≥ 1.0 Å from the DSRP group **(Fig. 5e)**. For the MIX group, there were more ensembles with lower global ΔRMSF predicted by the multi-conformation approach for all peptides in the group, but fewer with the RMSF_NMR_ ≥ 1.0 Å criteria applied. Collectively, these analyses suggest that the multi-conformation AF2 approach generates predictions with closer resemblance to NMR coordinate variability for DSRP peptides with RMSF_NMR_ ≥ 1.0 Å, whereas the conventional AF2 approach results in predictions with closer resemblance for AH MP peptides.

**Fig. 5:**
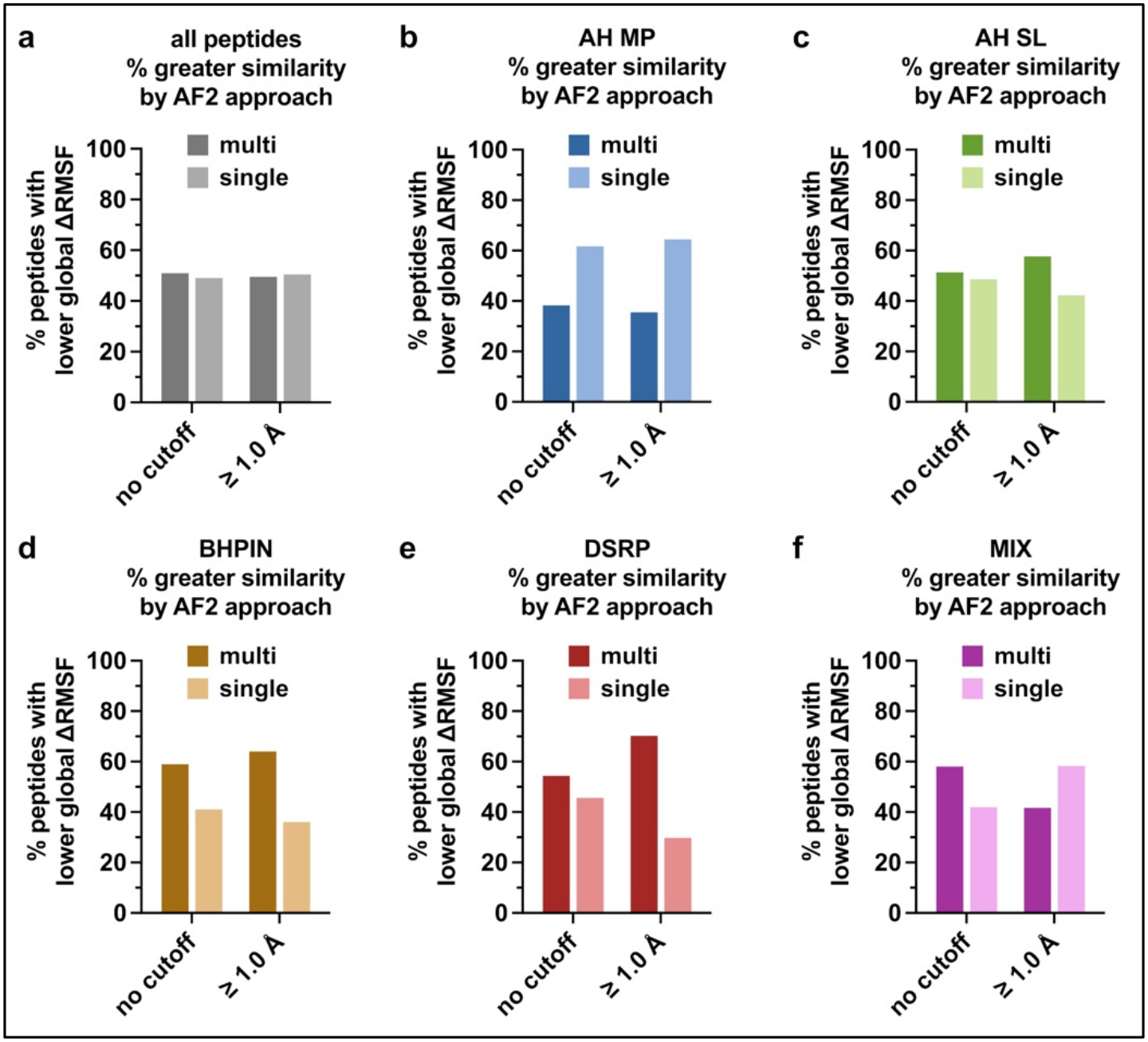
Percentage of greater experimental similarity by AF2 approach. **(a-f)** Percentage of peptides for which the global ΔRMSF between AF2 and NMR ensembles is lower for either the multi-conformation AF2 approach using multiple seeds and MSA subsampling (multi) or the conventional AF2 approach using single seeds without subsampling (single). Percentages are shown for peptides across the entire dataset (a) and among the different peptide groupings (b-f) with either no RMSF_NMR_ exclusion criteria (no cutoff) or including only peptides with RMSF_NMR_ ≥ 1.0 Å.

### Case study on the sarafotoxin-B peptide

Sarafotoxin-B, the DSRP peptide with the maximum composite score (PDB 1SRB) [33] **(Fig. 3c and d)**, was selected for a biologically-relevant case study. Sarafotoxin-B, also known as S6B, is a cardiotoxic peptide found in the venom of *Atractaspis engaddensis* snake [34]. Mechanistically, it binds to the endothelin receptor B and causes potentially lethal vasoconstriction [33, 35, 36]. The structure of sarafotoxin-b in complex with endothelin receptor B has recently been characterized by X-ray crystallography (PDB 6LRY) [35]. Given that peptides have the propensity to change conformation upon receptor binding, we compared how the NMR-determined and AF2-predicted conformers of unbound sarafotoxin-B differed from the endothelin receptor B-bound form of the peptide. Notably, the multi-conformation AF2 approach predicted conformers with lower RMSD, *i*.*e*., greater similarity, than NMR conformers to the receptor-bound form of sarafotoxin-b from the crystal structure **(Fig. 6a and b)**. This is not entirely surprising considering the inclusion of crystal structures in AF2 training [1], yet it does highlight the conformational diversity of the multi-conformation AF2 approach. Furthermore, while predictions of the multi-conformation AF2 approach and the conventional AF2 approach were not significantly different overall in terms of RMSD relative to the receptor-bound form of sarafotoxin-b, the multi-conformation AF2 approach predicted conformers with a broader RMSD range, revealing certain conformations with lower RMSD than the conventional AF2 approach predictions **(Fig. 6b)**. These observations highlight the ability of the multi-conformation AF2 approach to generate a broader range of peptide conformations that may provide additional insights into peptide structural variability in the context of biologically relevant receptor interactions.

**Fig. 6:**
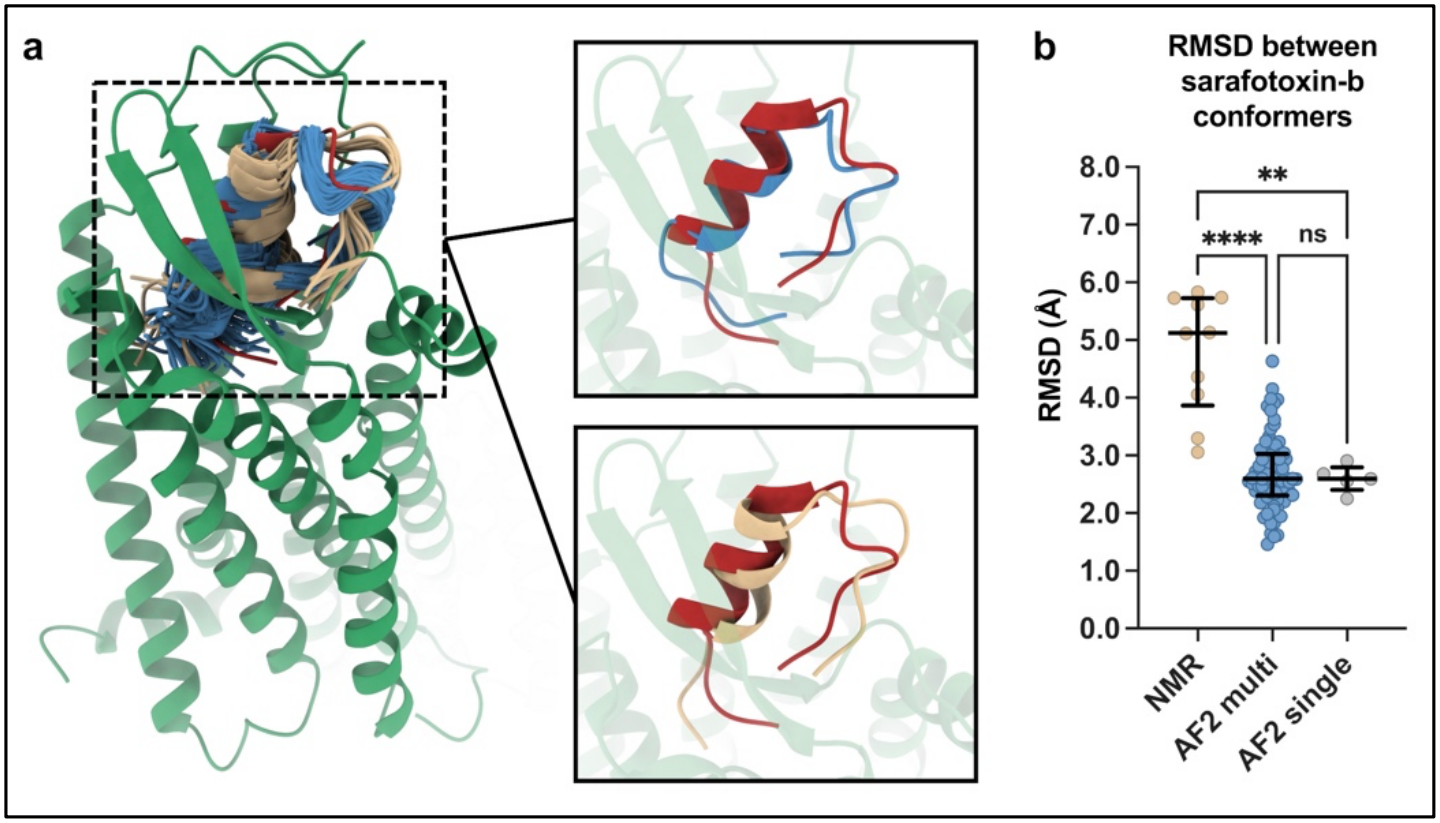
Experimental and predicted conformations of sarafotoxin-B in reference to its receptor-bound state. **(a)** Crystal structure of sarafotoxin-B (red) bound to endothelin receptor B (green) (PDB 6LRY) with overlay of the NMR-determined (beige) and multi-conformation AF2-predicted (blue) conformers. Conformers with the lowest RMSD relative to the receptor-bound form of sarafotoxin-B are depicted on the right: the most similar multi-conformation AF2-predicted conformer (upper box), RMSD = 1.4618 Å; and the most similar NMR-determined conformer (lower box), RMSD = 3.0560 Å. **(b)** RMSD between receptor-bound sarafotoxin-B and the unbound conformers from NMR, multi-conformation AF2 prediction (AF2 multi), and conventional AF2 prediction (AF2 single). Data represented as medians and corresponding IQRs. Statistical analysis by Kruskal Wallis one-way ANOVA with post-hoc Dunn’s multiple comparisons test, \*\*\*\**p* < 0.0001 (NMR vs. AF2 multi), \*\**p* = 0.0071 (NMR vs. AF2 single), ns (not significant, *p* ≥ 0.05).

## DISCUSSION

In the current work, we utilized AF2 for high-throughput prediction of peptide conformational ensembles using multiple seeds and MSA subsampling. This was applied to predict structural conformations of a dataset of 557 peptides categorized into five distinct structural groups, originating from a curated peptide dataset with NMR-determined conformational ensemble coordinates previously used for conventional AF2-based prediction [16]. The study from which the peptide dataset originates employed the conventional AF2 prediction approach (single seed without MSA subsampling) which generates five structural models per peptide [16], whereas the adapted multi-conformation approach [10-13] which we employed generates ensembles consisting of 80 structures per peptide. The two approaches differ in scope and intended use, with the conventional AF2 approach intended for static structure prediction and the multi-conformation AF2 approach intended for predicting conformational ensembles [1, 12]. As such, the present work should not be interpreted as a competitive comparison with the results reported by McDonald et al. [16], but rather as a complementary investigation pursuing a distinct direction of inquiry. It should also be noted that NMR ensemble coordinate data originate from study-dependent experimental conditions and differing conformer selection criteria, resulting in non-uniform representations of conformational variability and thereby posing a limitation on their use as references for comparisons with predicted ensembles [31, 32].

Following *de novo* computation of conformational ensembles of the 557 peptides, our analyses compared the structurally distinct groups of AH MP, AH SL, BHPIN, DSRP, and MIX with respect to co-evolutionary sequence (MSA depth) and structure (structured region RMSD and global ΔRMSF). The structured region RMSD and global ΔRMSF metrics served to measure predicted *versus* experimental similarity for rigid structure and conformational variability, respectively, with NMR ensemble coordinates used as experimental references. Taken together, our results indicate that the multi-conformation AF2 approach produced predictions that showed closer agreement with NMR ensemble coordinates for disulfide-rich peptides than for α-helical peptides, particularly membrane-associated α-helical peptides. It may also be the case that disulfide-rich peptides are inherently simpler prediction targets, given that disulfide bonds are typically expected to constrain flexibility, although it should also be considered that disulfide bonds do not uniformly affect conformational variability and their implications in this regard are complex [37-39].

Individual MSA depth comparisons among the peptide groups revealed one significant difference: MSA depth was greater for the DSRP group than for the AH MP group **(Fig. 1e)**. The DSRP group also had a weak but significant negative correlation between MSA depth and global ΔRMSF, *i*.*e*., greater MSA depth correlated with greater similarity in conformational variability between predicted and NMR ensemble coordinates **(Fig. 2e)**. Paradoxically, the opposite was observed for peptides from the α-helical AH MP and AH SL peptide groups **(Fig. 2b and c)**. This finding is intriguing considering the greater MSA depth of the DSRP group compared to the AH MP group, and may indicate that co-evolutionary sequence information is relevant, to some extent, for predicting conformational variability in addition to predicting static structure. In terms of structured region RMSD, significant differences were only found for the MIX group which had significantly greater RMSD compared to all other groups **(Fig. 1f)**. This difference may reflect the structurally heterogeneous nature of the MIX group, which contains both membrane-associated and soluble peptides with mixed secondary structure.

Although structured region RMSD comparisons did not indicate differences between the disulfide-rich and α-helical peptides, significant differences between these peptides were found among global ΔRMSF comparisons. Specifically, the DSRP group exhibited significantly lower global ΔRMSF than the AH MP and AH SL groups **(Fig. 1g)**, coinciding with the MSA depth and correlation observations for the DSRP and AH MP groups **(Fig. 1e; Fig. 2b and e)**, as well as the underestimation of flexibility among predicted ensembles of the AH MP and AH SL groups **(Fig. S3a and b)**. The greater similarity between AF2 and NMR ensemble coordinates of disulfide-rich peptides compared to α-helical peptides may, in part, be influenced by co-evolutionary sequence information available from the differing MSA depths. The paradoxical correlation between increasing MSA depth and increasing global ΔRMSF for peptides of the AH MP and AH SL groups **(Fig. 2b and c)** along with underestimation of flexibility in the same groups **(Fig. S3a and b)** may indicate that prediction of conformational variability for α-helical peptides is inherently challenging for AF2.

Comparing the multi-conformation AF2 approach and conventional AF2 approach provided additional insights into how well predictions by each approach resemble NMR reference coordinates. The finding that predictions by the multi-conformation approach for the DSRP group of peptides exhibit greater similarity to corresponding NMR ensemble coordinates, particularly for peptides with RMSF_NMR_ ≥ 1.0 Å, complements the other favorable observations made of the DSRP group. Likewise, the closer resemblance to NMR ensemble coordinates found for predictions of peptides from the AH MP group by the conventional approach further highlights the observed limitations of the multi-conformation approach in handling α-helical peptides. The multi-conformation approach seems to have performed favorably for predicting ensembles of BHPIN peptides based on the categorical percentage analysis. Although the differences between predictions generated by the two approaches were relatively modest overall, the multi-conformation approach carries the added benefit of computing 80 different structures compared to only five structures with the conventional approach. As such, the larger number of structures generated by the multi-conformation approach may reveal specific conformational configurations that may not appear with the conventional approach, although whether these additional conformations correspond to experimentally populated states was not evaluated here.

Given the nature of the peptide dataset and the parameterization of the modeling approach, several limitations should be considered. In terms of the NMR ensemble coordinates used as references, structures deposited in the PDB originate from study-dependent experimental methods and differing conformer selection criteria, introducing inconsistencies in the representation of conformational variability that may affect agreement with predicted ensembles. NMR observables may also be informative for assessing conformational variability. However, such data are not consistently available and vary in format, completeness, and quality, making them difficult to incorporate into large-scale analysis. The RMSD and RMSF metrics used in the present study assess agreement between predicted and experimentally determined structural ensemble coordinates rather than agreement with underlying NMR observables. Therefore, the present analyses should not be interpreted as validation against all forms of experimental NMR data, some of which may provide complementary insights for further assessing prediction accuracy. Furthermore, the multi-conformation prediction approach is computationally demanding, and together with the large scale of the dataset necessitated a standardized set of prediction parameters [12] (as outlined in the Methods section). Specific refinements of prediction parameters for select peptides may lead to better agreement with experimental references but are not scalable to this dataset. Future work would benefit from examining a smaller number of select peptides with NMR ensembles generated using uniform experimental methodologies for incorporation of NMR observables into analysis, and from evaluating multiple prediction parameter settings.

In conclusion, AF2-based multi-conformation prediction with multiple seeds and MSA subsampling offers a unique *in silico* approach for the study of peptide ensembles. This approach showed comparatively closer agreement with NMR ensemble coordinates for disulfide-rich peptides, with some favorable agreement also observed for β-hairpin-containing peptides. However, α-helical peptides present a significant challenge and reveal confounding MSA correlations. Peptides with mixed secondary structure are also less favorable candidates for this approach. Moreover, varying levels of inconsistency with NMR reference ensembles are evident across all peptide types that were assessed. Predictions should therefore be interpreted with discretion and an expectation for discrepancies relative to experimental data. Further research is needed to advance the current state of this multi-conformation prediction approach as it applies to peptides, and experimenting with differing AI/ML model architectures, incorporating specialized training regimens which include experimental peptide conformational ensembles, and alternative modes of MSA utilization may serve as effective strategies for improvement.

## METHODS

### AF2 Predictions

Predictions were performed using the ColabFold [12] implementation of AF2 in a Google Colab Jupyter Notebook environment with Hardware Accelerator set to T4 GPU. For the multi-conformation AF2 approach, the ColabFold conformation prediction protocol was used, as previously described [12]. This involved adjustment of the num_seeds parameter to 16 and adjustment of the max_msa parameter to 16:32. The original protocol calls for max_msa to be set to 32:64 for a larger protein [12], which was reduced to 16:32 in the current work in consideration of the shorter sequence length of peptides compared to proteins. All other parameters were left unmodified from their default settings, as follows:

- num_relax: 0
- template_mode: none
- msa_mode: mmseqs2_uniref_env
- model_type: auto
- num_recycles: 3
- recycle_early_stop_tolerance: auto
- use_dropout: unselected

By default, AlphaFold provides five structural models per individual prediction. As such, five structures were computed per seed, and with num_seeds set to 16 a total of *n* = 80 structures per peptide. In total, 44,560 individual structures were predicted across the entire set of 557 peptides.

For the conventional AF2 approach, the ColabFold monomer prediction protocol was used [12], with parameters left unmodified from their default settings, as follows:

- num_seeds: 1
- max_msa: auto
- num_relax: 0
- template_mode: none
- msa_mode: mmseqs2_uniref_env
- model_type: auto
- num_recycles: 3
- recycle_early_stop_tolerance: auto
- use_dropout: unselected

With five structures computed per seed, a total of 2,785 individual structures were predicted across the full set of 557 peptides using the conventional AF2 approach.

### Structural Analysis

RMSD and RMSF values were computed using custom implementations of Python code available online from the CHARMM-GUI lecture series [40, 41]. Structure alignment for RMSD calculations was restricted to Cα atoms of structured regions and RMSD calculations included only Cα atoms of these structured regions, as annotated in the original peptide dataset by McDonald et al. [16]. The coordinates of each structure in a given AF2-predicted conformational ensemble were compared to each structure in the corresponding NMR-determined conformational ensemble **(Fig. S1a)**. The average of all individual structure comparisons between one AF2 ensemble and the corresponding NMR ensemble provided the structured region RMSD value for each individual peptide. Subsequently, the structured region RMSD averaged across all peptides was calculated using the averaged RMSD values for each individual peptide. It should be noted that, because of the intrinsic structural heterogeneity within any given conformational ensemble, the RMSD within the ensemble holds a non-zero value. In other words, when comparing identical ensembles or comparing each structure within an ensemble to every other structure within the same ensemble, there is an inherent non-zero RMSD. Therefore, when comparing two non-identical ensembles, i.e., an AF2-predicted ensemble versus an NMR-determined ensemble, the calculated RMSD contains both ‘within-ensemble’ variability as well as ‘between-ensemble’ variability.

For RMSF determination, alignments involved the coordinates of all Cα atoms and were not restricted to structured regions. All structures in an ensemble were first aligned to an initial reference structure (i.e., the first structure in the ensemble) to compute the ensemble mean structure, the structure with the lowest RMSD relative to this mean was selected as the centroid, and all structures were then realigned to the centroid prior to RMSF calculation. Per-residue RMSF values were calculated based on the coordinates of Cα atoms for each residue across all structures within a given conformational ensemble. Per-residue RMSF values were used to calculate the global ΔRMSF metric (Equation 1) between coordinates of the two corresponding conformational ensembles, AF2-predicted and NMR-determined, for each peptide **(Fig. S1b)**. The global ΔRMSF metric first considers the absolute value of the difference between the RMSF of each individual residue pair (AF2-predicted versus NMR-determined) and then takes the average of the summed per-residue differences to provide a ‘global’ (overall) RMSF comparison between the two ensembles. Taking the absolute value of each per-residue RMSF difference is necessary in order to account for cancellation of positive and negative differences, which would otherwise skew the average of the summed differences towards 0 Å and thereby artificially overestimate accuracy.

### Statistical Analyses and Graphical Representation

Statistical analyses of IQRs, Kruskal Wallis one-way ANOVA, Dunn’s multiple comparisons test, Spearman correlation, nonlinear regression, and Wilcoxon matched-pairs test, were performed using GraphPad Prism 10. Statistical significance was set at *p* < 0.05. Categorical percentages were determined using Microsoft Excel. Graphical representations of data shown in **Fig. 1-5** and **Fig. S2-S9** were prepared using GraphPad Prism 10. Data used for statistical analyses are presented in **Supplementary Tables 1, 2, and 3**.

## ACKNOWLEDGEMENTS

This work was supported in part by the Levy-Longenbaugh Donor-Advised Fund to W.A. and R.P. RCSB Protein Data Bank is jointly funded by the National Science Foundation (DBI-2321666, PI: S.K.B.), the US Department of Energy (DE-SC0019749, PI: S.K.B.), and the National Cancer Institute, the National Institute of Allergy and Infectious Diseases, and the National Institute of General Medical Sciences of the National Institutes of Health (R01GM157729, PI: S.K.B.). This work was also supported by the National Science Foundation (PHY-2210291, PI: J.N.O.), National Institutes of Health (R35GM153502-01, PI: P.C.W.) and the Welch Foundation (C-1792, to J.N.O.). Work at the Center for Theoretical Biological Physics was supported by the National Science Foundation (PHY-2019745). J.N.O. is a Cancer Prevention Research in Texas Scholar in Cancer Research.

The annotated peptide dataset curated by McDonald et al. [16] was a foundational aspect of this work. AF2 predictions were performed by using ColabFold, an open-source platform developed by the research groups of Dr. Sergey Ovchinnikov at Massachusetts Institute of Technology and of Dr. Martin Steinegger at Seoul National University. Molecular analyses were performed with custom implementations of Python code available online from the CHARMM-GUI lecture series developed by the research group of Dr. Wonpil Im at Lehigh University. Molecular graphics were visualized with UCSF ChimeraX, developed by the Resource for Biocomputing, Visualization, and Informatics at the University of California, San Francisco, with support from the National Institutes of Health R01-GM129325 and the Office of Cyber Infrastructure and Computational Biology, National Institute of Allergy and Infectious Diseases [42]. The NMR ensemble depicted in the graphical abstract was obtained from the Protein Data Bank (PDB 2K2Y).

## AUTHOR CONTRIBUTIONS

A.M.I., C.M., S.K.B., P.C.W., J.N.O., W.A., and R.P. conceptualization; A.M.I. and C.M. methodology; A.M.I. investigation; A.M.I., C.M., S.K.B., P.C.W., J.N.O., W.A., and R.P. formal analysis; A.M.I. and C.M. writing—original draft; S.K.B., P.C.W., J.N.O., W.A., and R.P. writing—review & editing; S.K.B., P.C.W., J.N.O., W.A., and R.P. funding acquisition; W.A. and R.P. overall project supervision.

