## Supplementary Information for "High-throughput prediction of peptide structural conformations"

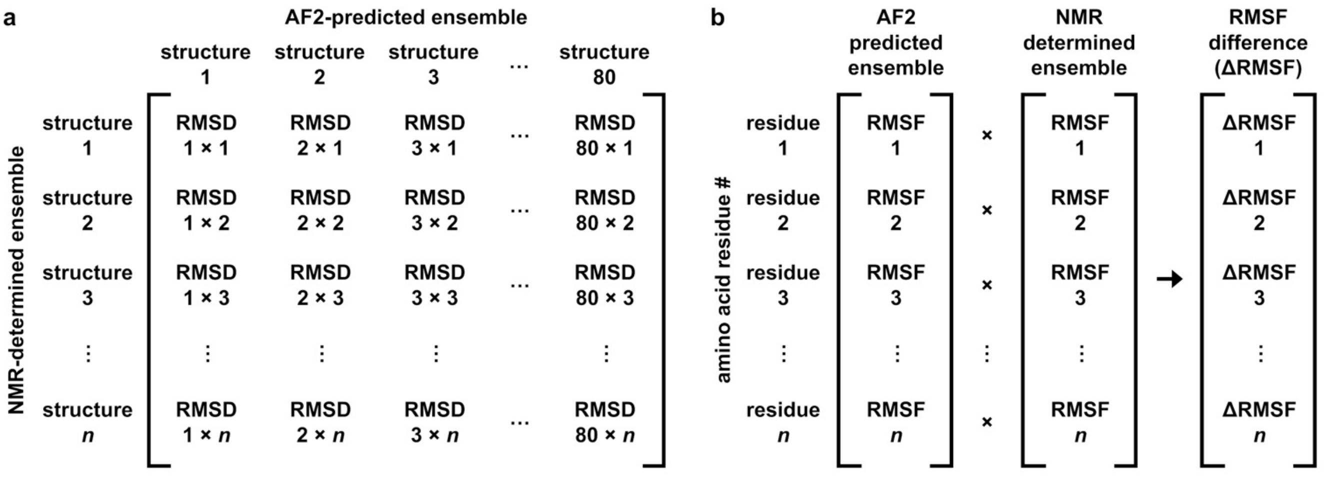


$$\textit{structured region RMSD} = \frac{\sum_{i=1}^{80n_{NMR}} RMSD_i}{80n_{NMR}}$$

$$\textit{global } \Delta RMSF = \frac{\sum_{i=1}^{n_r} |\Delta RMSF_i|}{n_r}$$

**Fig. S1: Overview of structural analysis metrics. (a)** Structured region RMSD was calculated by comparing each of the 80 structures from a given AF2-predicted conformational ensemble with all of the structures in the corresponding NMR-determined conformational ensemble, $n_{NMR}$, for a total of 80 $n_{NMR}$ comparisons. Only residues within structured regions were included, as outlined for each peptide in Table S1. **(b)** Global ΔRMSF was calculated by first determining the RMSF for each residue within a given AF2-predicted conformational ensemble and then within the corresponding NMR-determined ensemble independently for all residues in the peptide, $n_r$. The sum of the RMSF difference (ΔRMSF) for each residue between the two ensembles was then divided by the total number of residues, $n_r$. The absolute value of each ΔRMSF was taken in order to account for cancellation of positive and negative RMSF differences.

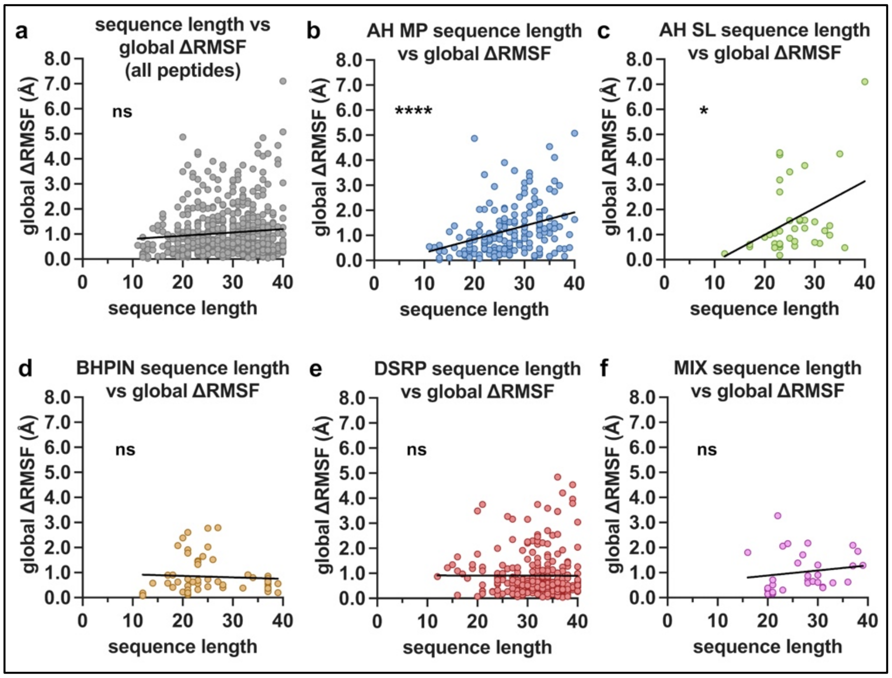


**Fig. S2: Associations of sequence length with global ΔRMSF between AF2-predicted and NMR-determined structural ensembles. (a-f)** Sequence length versus Cα global ΔRMSF for all peptides **(a)** and for the different peptide groups **(b-f)**. Statistical analyses by Spearman correlation: **(a)** all peptides $r$ = 0.03304, 95% CI -0.05263 to 0.1182, ns (not significant, $p \geq 0.05$), $n$ = 557 peptides; **(b)** AH MP $r$ = 0.3570, 95% CI 0.2148 to 0.4844, ****$p$ = < 0.0001, $n$ = 172 peptides; **(c)** AH SL $r$ = 0.3724, 95% CI 0.04505 to 0.6275, *$p$ = 0.0232, $n$ = 37 peptides; **(d)** BHPIN $r$ = 0.1025, 95% CI -0.1726 to 0.3627, ns, $n$ = 56 peptides; **(e)** DSRP $r$ = -0.06590, 95% CI -0.1893 to 0.05957, ns, $n$ = 261 peptides; **(f)** MIX $r$ = 0.3347, 95% CI -0.03320 to 0.6227, ns, $n$ = 31 peptides. For all panels, linear regression trend lines are depicted in black.

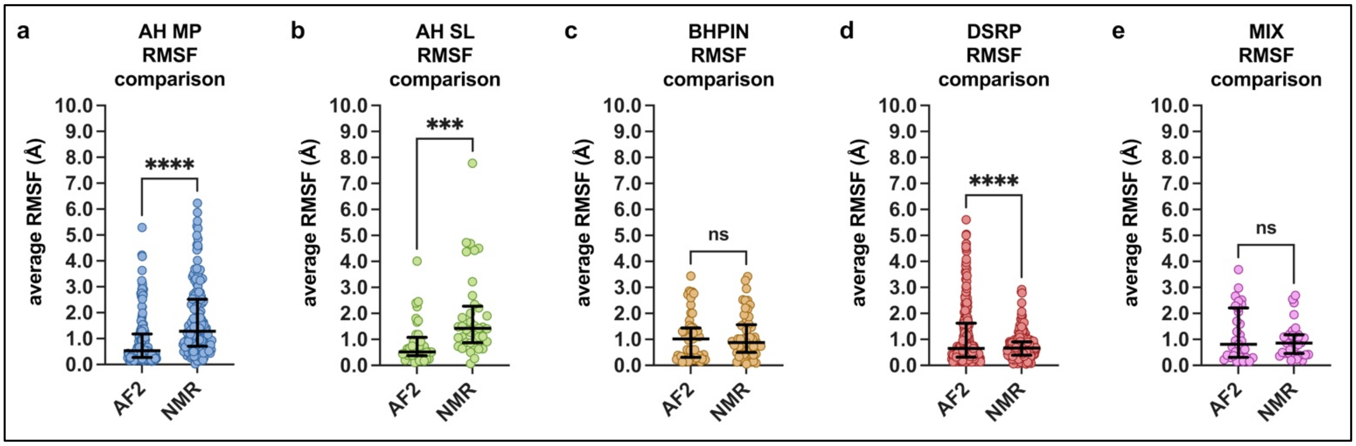


**Fig. S3: Average RMSF values of individual conformational ensembles. (a-e)** Comparisons of average RMSF values of individual AF2 and NMR ensembles. Data represented as medians and corresponding IQRs. Statistical analysis by Wilcoxon matched-pairs test, ***$p = 0.0006$ (AF2 median = 0.5154 Å, NMR median = 1.4246 Å), AH MP ****$p < 0.0001$ (AF2 median = 0.5268 Å, NMR median = 1.2844 Å), DSRP ****$p < 0.0001$ (AF2 median = 0.6511 Å, NMR median = 0.6651 Å).

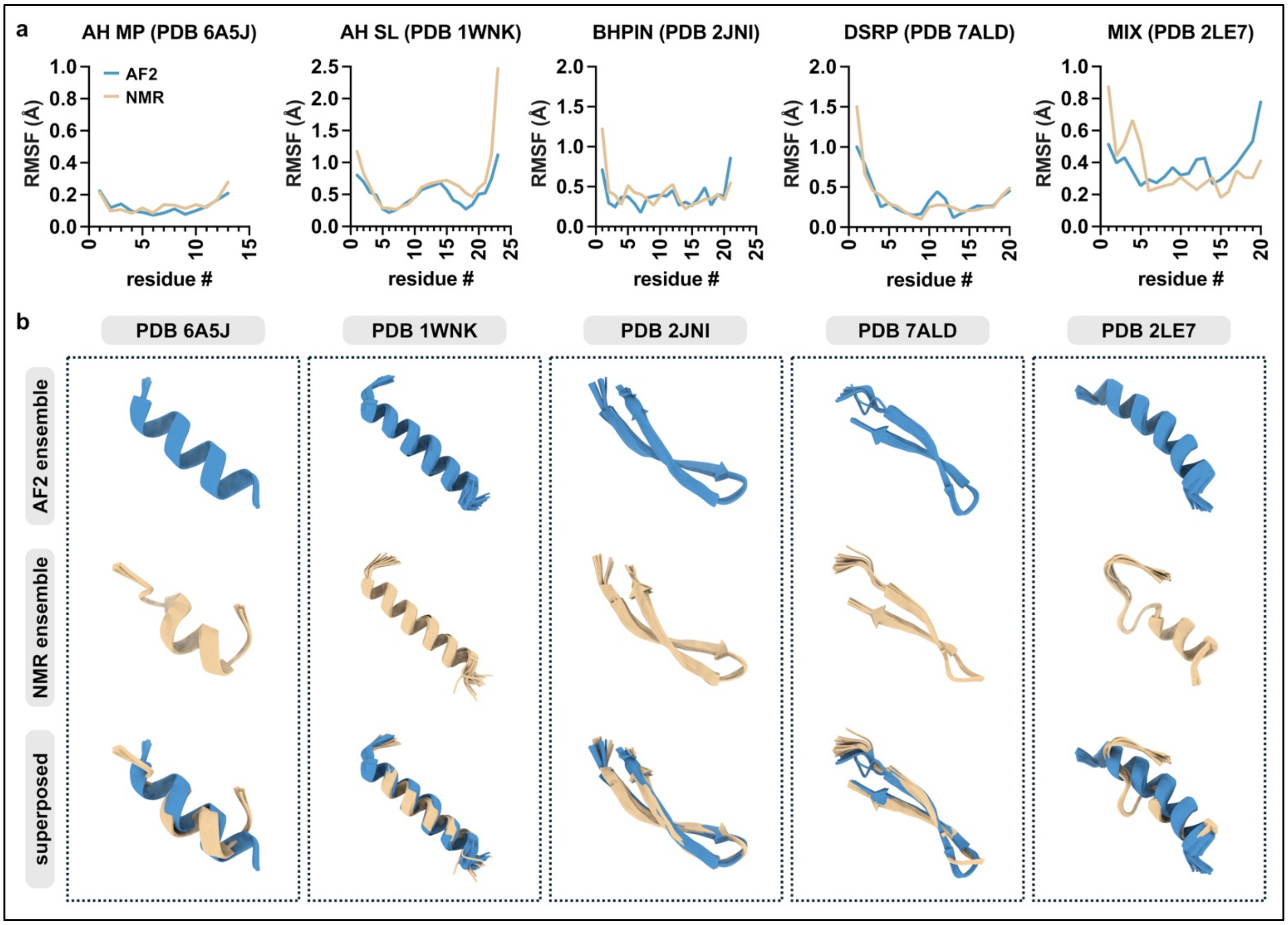


**Fig. S4: Examples of peptides with both low composite score and sub-angstrom conformational variability of experimental ensembles. (a)** Comparisons of Cα RMSF values of AF2-predicted and NMR-determined conformational ensembles with low composite score and sub-angstrom $RMSF_{NMR}$ from each peptide group. **(b)** AF2-determined (blue), NMR-determined (beige), and superposed structures of the representative conformational ensembles from (b). Global ΔRMSF values are as follows: PDB 6A5J, 0.0267 Å; PDB 1WNK, 0.1741 Å; PDB 2JNI, 0.1186 Å; PDB 7ALD, 0.0778 Å; and PDB 2LE7, 0.1413 Å.

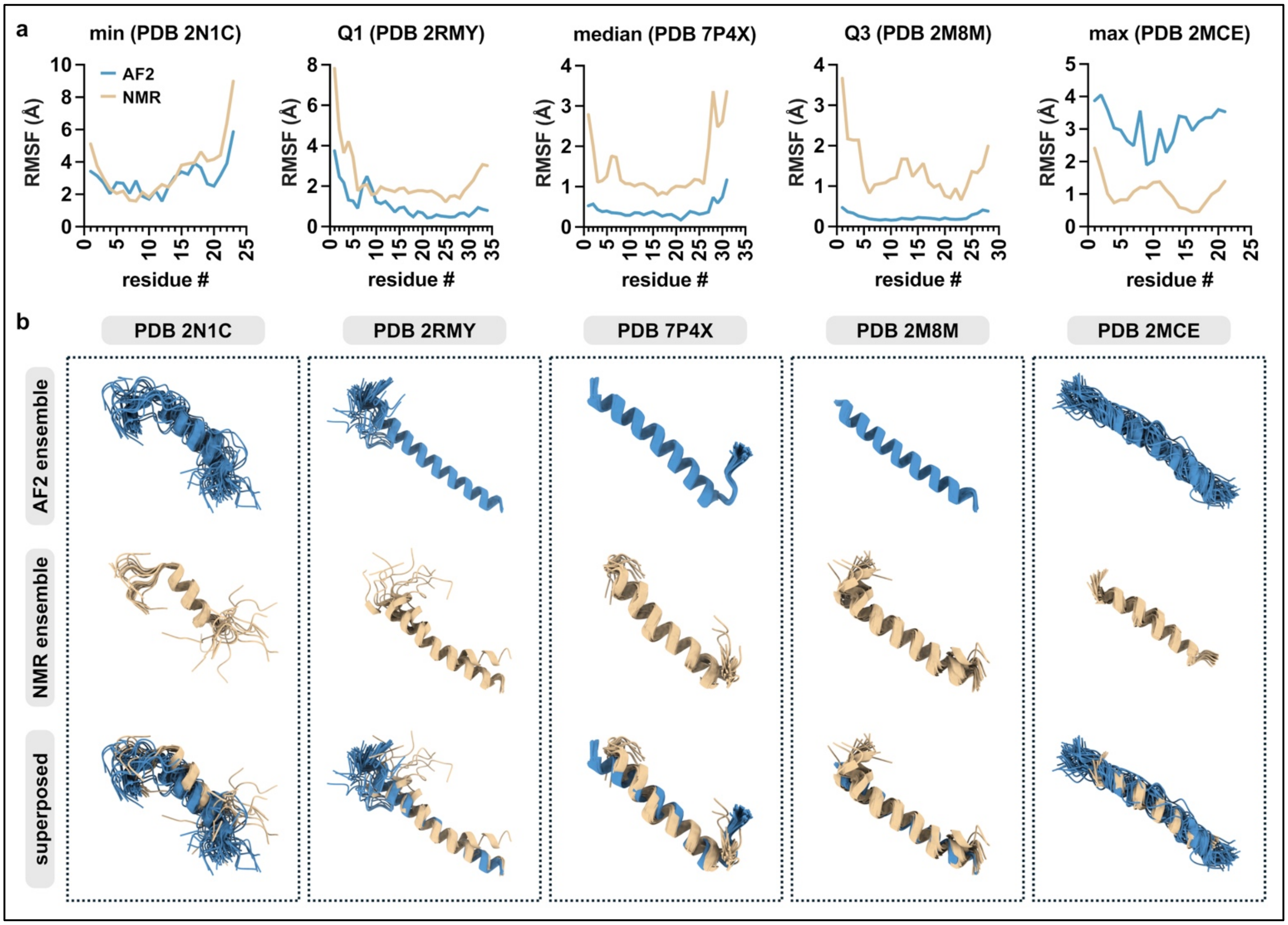


**Fig. S5: AH MP composite score quartile-representative peptide examples. (a)** Comparisons of Cα RMSF values of AF2-predicted and NMR-determined conformational ensembles representative of the minimum, Q1, median, Q3, and maximum composite scores from the AH MP peptide group for peptides with $RMSF_{NMR} \geq 1$ Å **(Fig. 3b)**. Representative ensembles were selected as the largest observed composite score ≤ each quartile. **(b)** AF2-determined (blue), NMR-determined (beige), and superposed structures of the representative conformational ensembles from (b). Global ΔRMSF values are as follows: PDB 2N1C, 0.8369 Å; PDB 2RMY, 1.2694 Å; PDB 7P4X, 1.0425 Å; PDB 2M8M, 1.1386 Å; and PDB 2MCE, 2.0344 Å.

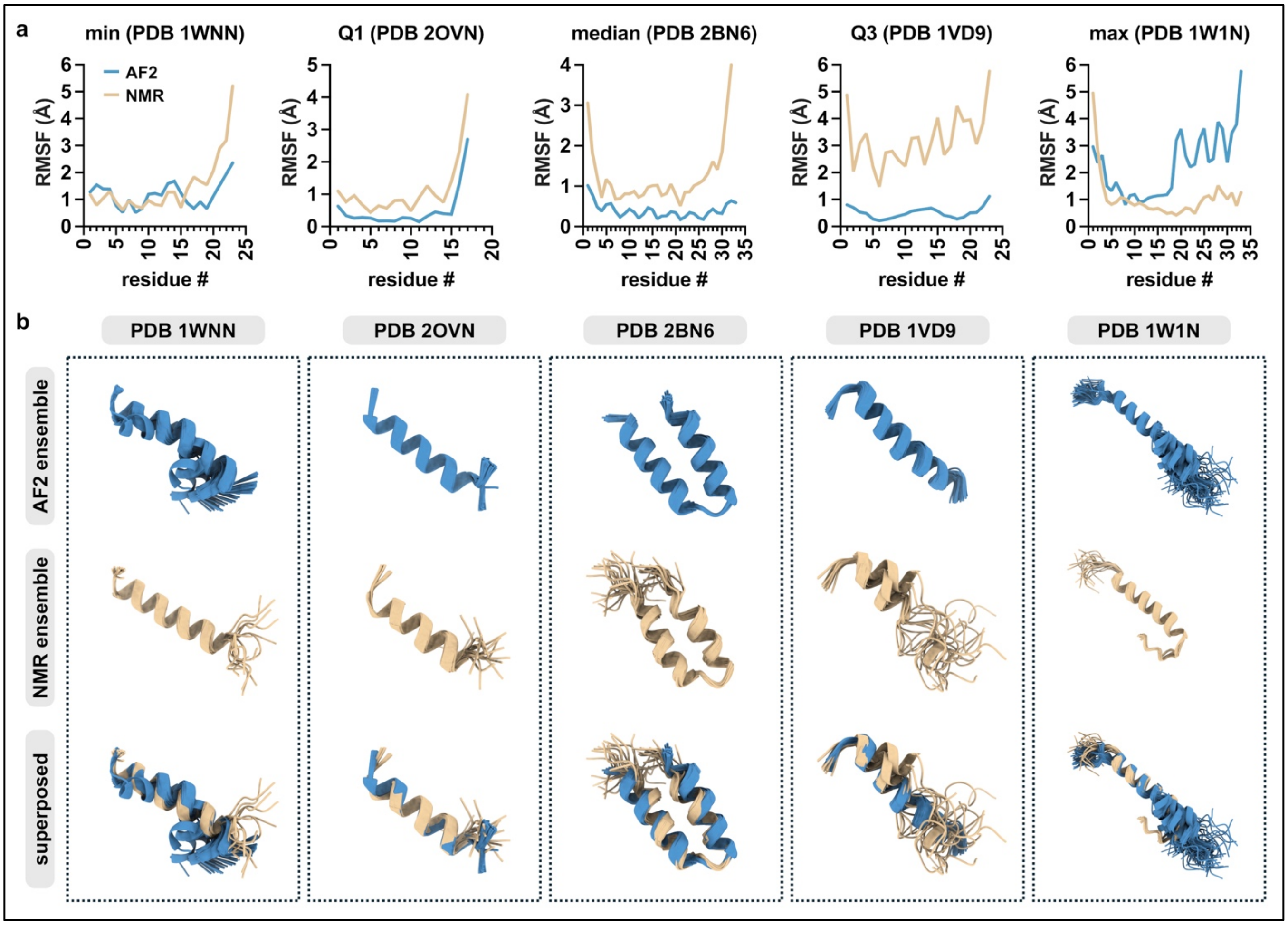


**Fig. S6: AH SL composite score quartile-representative peptide examples. (a)** Comparisons of Cα RMSF values of AF2-predicted and NMR-determined conformational ensembles representative of the minimum, Q1, median, Q3, and maximum composite scores from the AH SL peptide group for peptides with $RMSF_{NMR} \geq 1$ Å **(Fig. 3b)**. Representative ensembles were selected as the largest observed composite score ≤ each quartile. **(b)** AF2-determined (blue), NMR-determined (beige), and superposed structures of the representative conformational ensembles from (b). Global ΔRMSF values are as follows: PDB 1WNN, 0.6023 Å; PDB 2OVN, 0.6097 Å; PDB 2BN6, 0.9773 Å; PDB 1VD9, 2.7033 Å; and PDB 1W1N, 1.3573 Å.

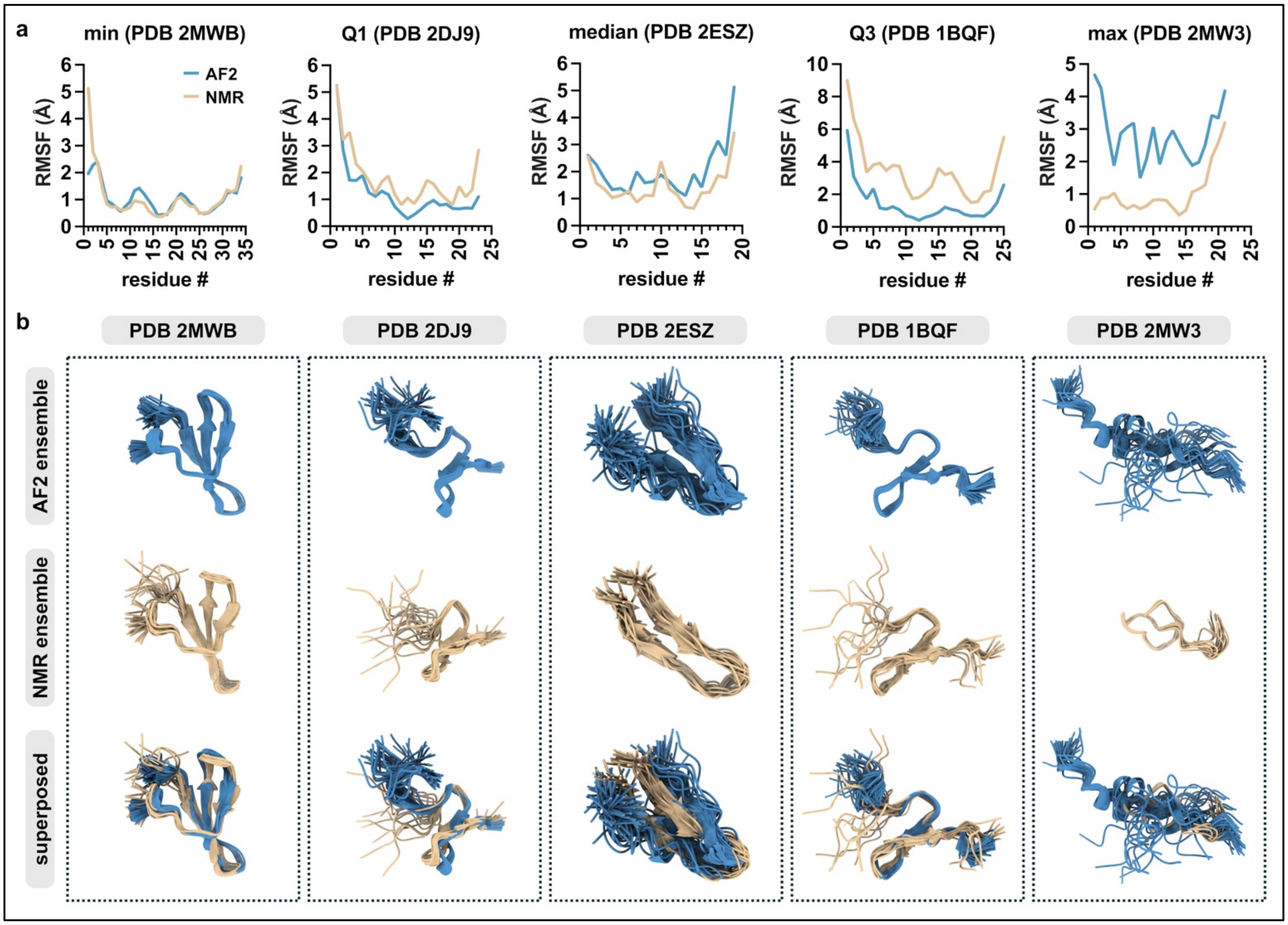


**Fig. S7: BHPIN composite score quartile-representative peptide examples. (a)** Comparisons of Cα RMSF values of AF2-predicted and NMR-determined conformational ensembles representative of the minimum, Q1, median, Q3, and maximum composite scores from the BHPIN peptide group for peptides with $RMSF_{NMR} \geq 1$ Å **(Fig. 3b)**. Representative ensembles were selected as the largest observed composite score ≤ each quartile. **(b)** AF2-determined (blue), NMR-determined (beige), and superposed structures of the representative conformational ensembles from (b). Global ΔRMSF values are as follows: PDB 2MWB, 0.2395 Å; PDB 2DJ9, 0.5650 Å; PDB 2ESZ, 0.6215 Å; PDB 1BQF, 2.0207 Å; and PDB 2MW3, 1.7763 Å.

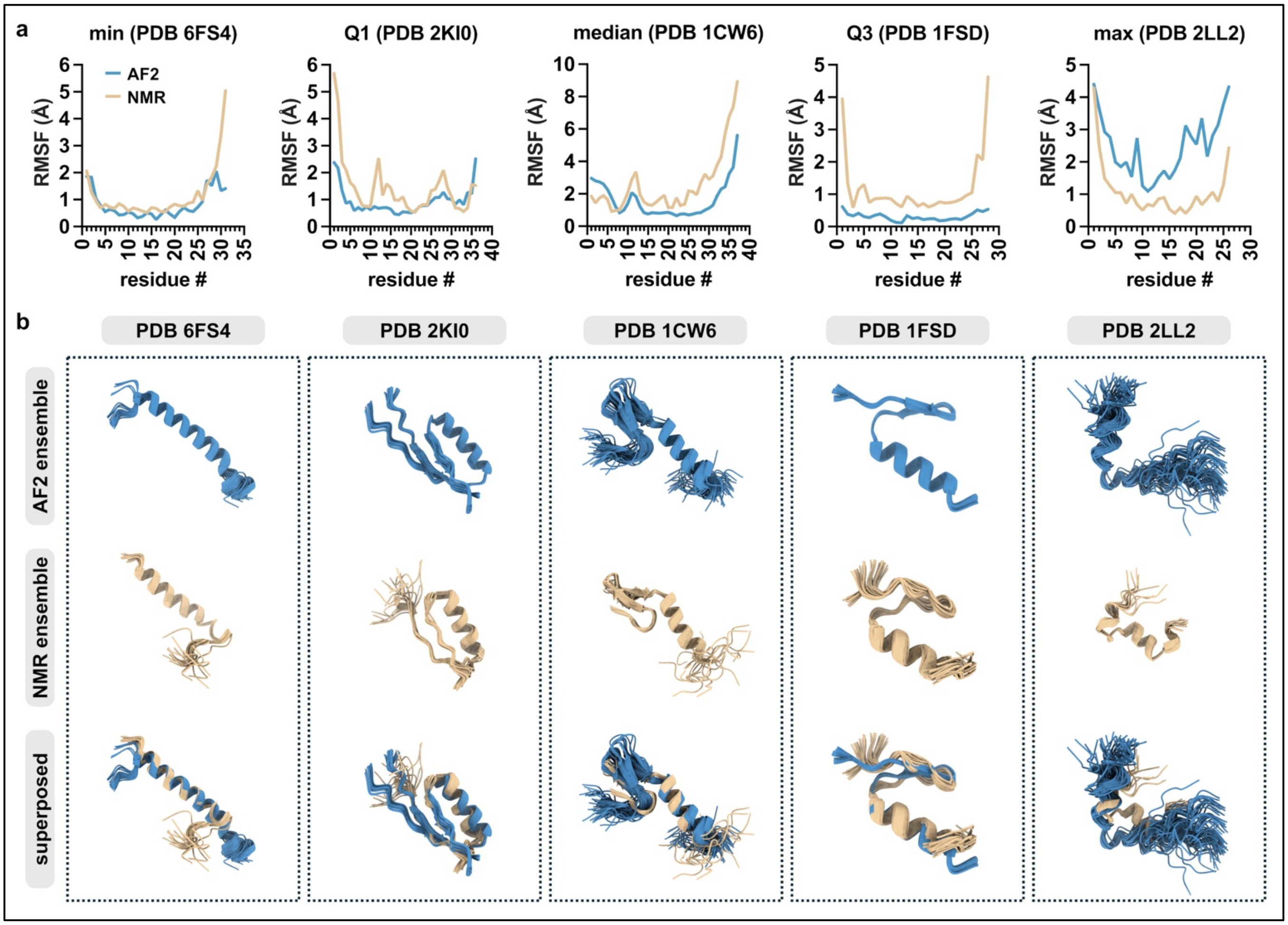


**Fig. S8: MIX composite score quartile-representative peptide examples. (a)** Comparisons of Cα RMSF values of AF2-predicted and NMR-determined conformational ensembles representative of the minimum, Q1, median, Q3, and maximum composite scores from the MIX peptide group for peptides with $RMSF_{NMR}$ ≥ 1 Å **(Fig. 3b)**. Representative ensembles were selected as the largest observed composite score ≤ each quartile. **(b)** AF2-determined (blue), NMR-determined (beige), and superposed structures of the representative conformational ensembles from (b). Global ΔRMSF values are as follows: PDB 6FS4, 0.3986 Å; PDB 2KI0, 0.6258 Å; PDB 1CW6, 1.2702 Å; PDB 1FSD, 0.8703 Å; and PDB 2LL2, 1.3804 Å.

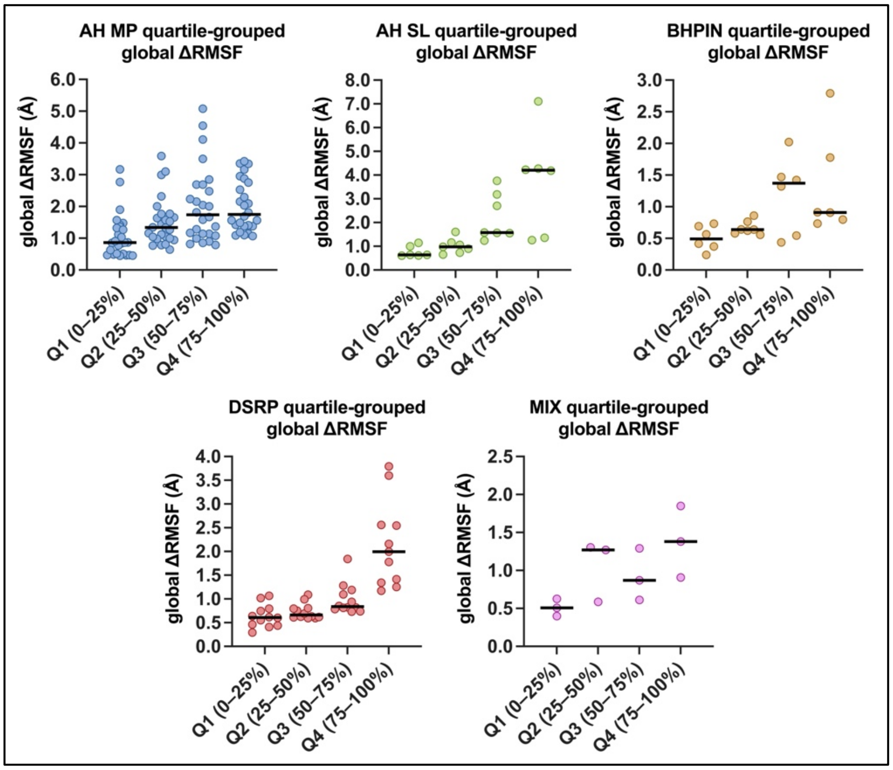


**Fig. S9: Composite score quartile groupings.** Global ΔRMSF between AF2 and NMR ensembles of each of the five peptide categories grouped by the composite score quartiles, *i.e.*, ranging between 0%, 25%, 50%, 75%, and 100% of the composite score distribution. Black lines represent medians. This analysis includes only peptides with $RMSF_{NMR}$ ≥ 1 Å, as outlined in Supplementary Table 2.
